# “Single-subject studies”-derived analyses unveil altered biomechanisms between very small cohorts: implications for rare diseases

**DOI:** 10.1101/2021.02.10.430623

**Authors:** Dillon Aberasturi, Nima Pouladi, Samir Rachid Zaim, Colleen Kenost, Joanne Berghout, Walter W. Piegorsch, Yves A. Lussier

**Affiliations:** Center for Biomedical Informatics and Biostatistics (CB2), University of Arizona, Tucson, AZ, USA; Dept. Of Medicine, University of Arizona, Tucson, AZ, USA; Graduate Interdisciplinary Prog. in Statistics & Data Science, University of Arizona, Tucson, AZ, USA; Ctr for Appl. Genetics and Genomic Medic., University of Arizona, Tucson, AZ, USA; Bio5 Institute; University of Arizona, Tucson, AZ, USA; Dept of Biomedical Informatics; University of Utah, UT, USA

## Abstract

**Motivation:** Identifying altered transcripts between very small human cohorts is particularly challenging and is compounded by the low accrual rate of human subjects in rare diseases or sub-stratified common disorders. Yet, <u>s</u>ingle-<u>s</u>ubject <u>s</u>tudies (S^3^) can compare paired transcriptome samples drawn from the same patient under two conditions (e.g., treated vs pre-treatment) and suggest patient-specific responsive biomechanisms based on the overrepresentation of functionally defined gene sets. These improve statistical power by: (i) reducing the total features tested and (ii) relaxing the requirement of within-cohort uniformity at the transcript level. We propose *Inter-N-of-1*, a novel method, to identify meaningful biomechanism differences between very small cohorts by using the effect size of “single-subject-study”-derived responsive biomechanisms.

**Results:** In each subject, *Inter-N-of-1* requires applying previously published S^3^-type *N-of-1-pathways MixEnrich* to two paired samples (e.g., diseased vs unaffected tissues) for determining patient-specific <u>e</u>nriched <u>g</u>enes <u>s</u>ets: Odds Ratios (S^3^-OR) and S^3^-variance using Gene Ontology Biological Processes. To evaluate small cohorts, we calculated the precision and recall of *Inter-N-of-1* and that of a control method (GLM+EGS) when comparing two cohorts of decreasing sizes (from 20 vs 20 to 2 vs 2) in a comprehensive six-parameter simulation and in a proof-of-concept clinical dataset. In simulations, the *Inter-N-of-1* median precision and recall are > 90% and >75% in cohorts of 3 vs 3 distinct subjects (regardless of the parameter values), whereas conventional methods outperform *Inter-N-of-1* at sample sizes 9 vs 9 and larger. Similar results were obtained in the clinical proof-of-concept dataset.

**Availability:** R software is available at Lussierlab.net/BSSD.

## 1 Introduction

Empirical evidence unveils a methodological gap when comparing transcriptomic differences in biomechanisms within very small human cohorts due to variations in heterogenicity, uncontrolled biology (age, gender, etc.), and diversity of environmental factors (nutrition, sleep, etc.). (Griggs, et al., 2009; Liu, et al., 2014; Schurch, et al., 2016; Soneson and Delorenzi, 2013). Paradoxically, rare diseases are common: 8% prevalence in the population (Elliott and Zurynski, 2015) and 26% of children who attend disability clinic (Guillem, et al., 2008). As timely and sizeable patient accrual of rare or micro-stratified diseases are prohibitive, there lies an opportunity for empowering clinical researchers with feasible statistical designs that enable smaller cohorts.

On the other hand, well-controlled isogenic studies (e.g., cellular models) can yield differentially expressed genes (DEGs) between two small samples. We and others have applied the power of the isogenic framework through the comparison of two sample transcriptomes from one subject in single-subject studies (S^3^). While *transcript-level* differences between two-sample remains inaccurate (Vitali, et al., 2017; Zaim, et al., 2019), *gene set-level* (*pathway/biosystem*) S^3^ have been shown to accurately discover altered biomechanisms from paired transcriptome samples drawn from the same patient under two conditions (e.g., tumor-normal, treated-untreated) (Ozturk, et al., 2018; Vitali, et al., 2017). The results of the S^3^ *gene set* analyses have been validated in various contexts such as cellular/tissular models (Balli, et al., 2019; Gardeux, et al., 2014; Gardeux, et al., 2015), retrospectively in predicting cancer survival (Li, et al., 2017; Schissler, et al., 2015; Schissler, et al., 2018) circulating tumor cells (Schissler, et al., 2016), biomarker discovery simulations (Zaim, et al., 2018), and therapeutic response (Li, et al., 2017). Despite the success of these models to derive effect sizes and statistical significance in single-subject studies of transcriptomes, these samples are isogenic or quasi-isogenic, and thus do not necessarily generalize to a group of subjects (*cohort-level signal*). To address the latter, we reported that determining single cohort-level significance by combining gene set signal (e.g., pathways) from S^3^ analyses can be more accurate than conventional DEG analyses followed by gene set enrichment analysis (GSEA) (Subramanian, et al., 2005) in small cohort simulations (Zaim, et al., 2018) and in previously published datasets (Li, et al., 2017)]. However, these methods still used simplistic cohort-level assumptions of centrality (median) and did not explore *comparing signal divergence between two cohorts*.

To address the methodological gap, we therefore hypothesized that single-subject transcriptomic studies of gene sets increase the transcriptomic signal-to-noise ratio within subject and lead to an improved signal between small patient cohorts, as small as 3vs3 subjects per group. While technically different from the analysis of the standard two factor interactions in conventional cohort statistics, the proposed framework is conceptually related to a statistical interaction in that a within-single-subject analysis (subject-specific transcriptome dynamics) is followed by within-group agreement for characterizing Factor 1 (e.g., cancer vs paired normal tissue) and between group comparisons (Factor 2; e.g., responsive vs unresponsive to therapy). The strategy improves the statistical power by: (i) reducing the total features tested (gene set-level rather than transcript-level), (ii) relaxing the requirement of within-cohort uniformity at the transcript level as the coordination is conducted at the gene set-level, and (iii) reducing confounding factors through the paired sample design of S^3^-analyses within subject. The novel bioinformatic method identifies meaningful biomechanism differences between very small cohorts by using single-subject-study-derived effect sizes for gene sets. Additionally, we show through both a simulation and a real data case example that within cohorts of varying sizes (3 to 7 subjects) this method outperforms traditional methods, which are based on generalized linear modeling followed by common gene set enrichment or overlap analysis. We then apply this novel method to the effect sizes of two different single-subject analyses to illustrate the flexibility and utility of the proposed method for a variety of inputs.

## 2 Methods

**Table 1** defines abbreviations and **Figure 1** provides an overview of the proposed new method (*Inter-N-of-1*). To motivate the development of transcriptome analytics between very small human samples, by nature heterogenicity, we first demonstrate the limitation of a Generalized Linear Model to DEGs between 23 TP53 and 19 PIK3CA breast cancer samples. Next, we describe two new methods *Inter-N-of-1* (MixEnrich) and *Inter-N-of-1* (NOISeq) and compare them to a Generalized Linear Model (implemented in *LIMMA*) (i) in simulation studies with parameters estimated from empirical analyses of real datasets and (ii) in a proof-of-concept study of breast cancer subjects. Also, the evaluation of the proposed new methods is conservative as it is conducted against a reference standard built with a distinct Generalized Linear Model (*edgeR*) using all samples.

**Fig. 1.**
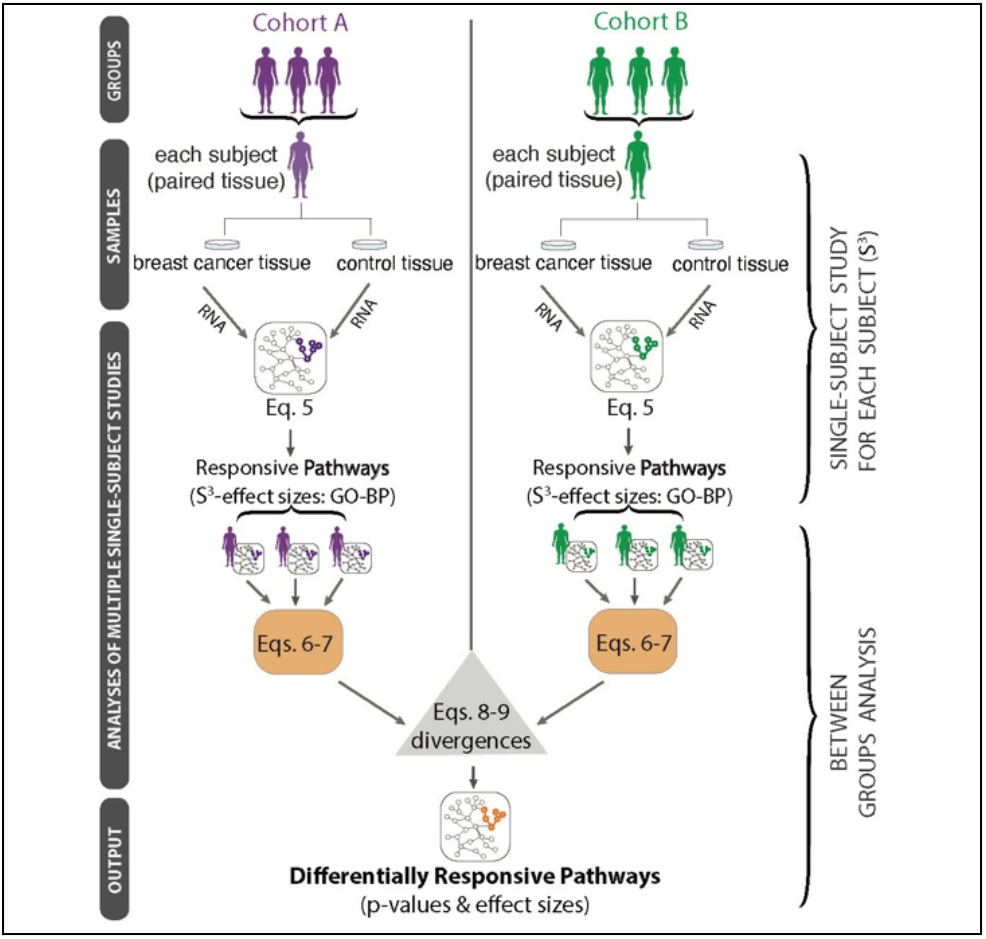
Overview of the gene set analyses (*Inter-N-of-1*) that leverage effect sizes and variances from single-subject studies to conduct subsequent group comparisons. Single-subject studies details are provided in **Figure 2**.

**Table 1.** Abbreviations

| Abbreviation | Term |
| --- | --- |
| DEG | Differentially Expressed Gene |
| Inter-N-of-1 | "Responsive Pathway Effect Size"-based cross cohort comparison |
| EGS | Enriched Gene Set of responsive pathways between two conditions within a single-subject-study (e.g., cancer vs control tissue) |
| FET | Fisher's Exact Test |
| FDR | False Discovery Rate |
| GEO | Gene Expression Omnibus |
| GLM+EGS | Generalized Linear Models with Enriched Gene Sets |
| GO-BP | Gene Ontology Biological Processes |
| GS | Gene Set (calculated from GO-BP) |
| ITS | Information Theory-Based Similarity between GO-BPs |
| log<sub>2</sub>FC | Log base 2 transformation of the transcripts fold-change |
| MLE | Maximum Likelihood Estimator |
| OR, S<sup>3</sup>-OR | Odds Ratio: S<sup>3</sup>-prioritized transcripts enriched in GO-BP |
| PCA | Principal Component Analysis |
| PIK3CA | Phosphatidylinositol-4,5-bisphosphate 3-kinase catalytic subunit alpha gene; HGNC:8975 |
| RSEM | RNA-Seq normalization by Expectation Maximization |
| S<sup>3</sup> | Single-Subject Studies |
| TP53 | Tumor protein p53 gene; HGNC:11998 |

### 2.1 Datasets

We obtained 5,179 g*ene sets* from Gene Ontology Biological Processes (GO-BP) (downloaded on 02/07/2019). For the determining realistic simulation parameters, we used two datasets (I and II) that are composed of paired samples.

(I) We downloaded 7 estrogen-stimulated and 7 unstimulated MCF7 breast cancer cells sample replicates provided by (Liu, et al., 2014) that were from the Gene Expression Omnibus (GEO) (Edgar, et al., 2002) on 10/14/2020. The sequences within the Sequence Read Archive files for the 30M reads of MCF7 cells were aligned using hg19 as the reference genome and the resulting RNA-seq counts were processed into fpkm units (Fragments Per Kilobase of transcript per Million mapped reads).

(II) We obtained 224 samples of paired breast cancer tumor and tissue-matched normal RNA-seq expression profiles (***Factor 1***) from the same subjects (n= 112) from The Cancer Genome Atlas (TCGA) Breast Invasive Carcinoma data collection (Cancer Genome Atlas, 2012; Ciriello, et al., 2015) using the Genomic Data Commons tools (Grossman, et al., 2016) (Obtained 10/22/2015). As a *proof-of-concept application* of the proposed methods, we sampled small groups of subjects from a subset of the TCGA breast cancer dataset comprising subjects with somatic (tumor) mutations in either *TP53* (n = 23) or *PIK3CA* (n = 19), but not both. *TP53* and *PIK3CA* (***Factor 2***) have been reported as the two most commonly mutated genes observed in breast cancer and differ as follows: (i) in expression patterns (Cancer Genome Atlas, 2012), (ii) cancer subtypes (Van Keymeulen, et al., 2015), (iii) clinical outcomes (Kim, et al., 2017), and (iv) responsiveness to specific therapies (Andre, et al., 2019). These data were downloaded using the R package TCGA2STAT(n=42 cases; 84 files) (Wan, et al., 2016).

#### Data access and preparation

(A) For the single-subject studies, we applied a three-stage filtering of the transcripts in which - within each sample pair – (i) we removed all transcripts with mean expression less than 5 counts, (ii) found the union of all genes remaining amongst all pairs, and (iii) excluded all genes not present in the union of these two steps (17,923 genes remaining). We added 1 to expression counts to eliminate “zeros”.

(B) For the generalized linear model-based analyses, we applied a different filtering process to the raw data where we eliminated all the transcripts with 0 counts for each subject and then calculated the coefficient of variation (CV) for each transcript. We selected the transcripts with CVs within the top 70 percentile of those remaining (13,932 genes remaining).

### 2.2 Proposed S^3^-anchored Responsive Pathway Effect Size Methods for comparing very small human cohorts

The following paragraphs will develop the methodology by which we conduct single-subject studies prior to cross-cohort comparison to discover the effect size of responsive pathways in each subject and increase the features signal-to-noise ratio. **Table 2** summarizes the variables.

**Table 2.** Variable Definitions

| Variable | Definition |
| --- | --- |
| $g_{gs,k_j}$ | The number of DEGs within gene set $gs$ for subject $k_j$ in cohort $K$ |
| $g'_{gs,k_j}$ | The number of genes NOT differentially expressed in gene set $gs$ for subject $k_j$ in cohort $K$ |
| $h_{gs,k_j}$ | The number of DEGs NOT in gene set $gs$ for subject $k_j$ in cohort $K$ |
| $h'_{gs,k_j}$ | Number of genes neither differentially expressed nor in gene set $gs$ for subject $k_j$ in cohort $K$ |
| $N$ | Number of gene sets |
| $P(\cdot)$ | Probability of Event $(\cdot)$ occurring |
| $\tilde{p}_{gs,\Delta}$ | p-value for gene set $gs$ produced by the Inter-N-of-1 |
| $Q_{gs,k_j}$ | Continuity-corrected log $S^3$-OR corresponding to gene set $gs$ for subject $k_j$ in cohort $K$ |
| $\bar{Q}_{gs,k_j}$ | The mean continuity-corrected log $S^3$-OR of in gene set $gs$ for subject $k_j$ in cohort $K$ |
| $S_K$ | The number of subjects in a cohort $K$ (e.g. those with a *PIK3CA* or with *TP53* somatic mutation) |
| $\theta_K$ | Expected value of the continuity corrected log $S^3$-OR for the molecular-defined cohort $K$ |
| $\text{var}(Q_{gs,k_j})$ | Variance associated with continuity-corrected log $S^3$-OR corresponding to gene set $gs$ for subject $k_j$ in cohort $K$ |
| $W_{gs,\Delta}$ | The test statistic for the *Inter-N-of-1* for gene set $gs$ |
| $Z$ | A standard normal random variable |

#### Identification of overrepresented gene sets for each subject

As illustrated in **Panel A** of **Figure 2,** we applied to each of the tumor-normal pairs the N-of-1-*pathways* MixEnrich method that we had previously developed and validated (Berghout, et al., 2018; Li, et al., 2017; Zaim, et al., 2019). Briefly, this method models the absolute value of the log_2_ transformed fold change (FC) for each gene across the two paired transcriptomes being studied and uses a probabilistic Gaussian mixture to assign a posterior probability that the gene is differentially expressed between tumor and normal conditions. Within the simulation, prioritized transcripts were defined as those with a posterior probability of being differentially expressed higher than 0.99. Within the TCGA breast cancer dataset, said definition included having both a posterior probability of being differentially expressed higher than 0.99 and an absolute-valued log_2_FC higher than log_2_(1.2). Genes were assigned to gene sets using the Gene Ontology (Ashburner, et al., 2000) Biological Process (GO-BP) hierarchy, filtered to those terms with gene set size between 15-500 genes, with subsumption to maximize interpretability. These DEGs were used to determine the overrepresented, or enriched, gene sets of interest using a two-sided Fisher’s Exact Test (FET) (Fisher, 1935) with a False Discovery Rate (FDR) of 5%. The output of this analysis generated lists of gene sets, with each list representing a single subject’s tumor-normal pair and comprising GO-BP terms accompanied by contingency table counts which were used to calculate an odds ratio (S^3^-OR) as the effect size.

**Fig. 2.**
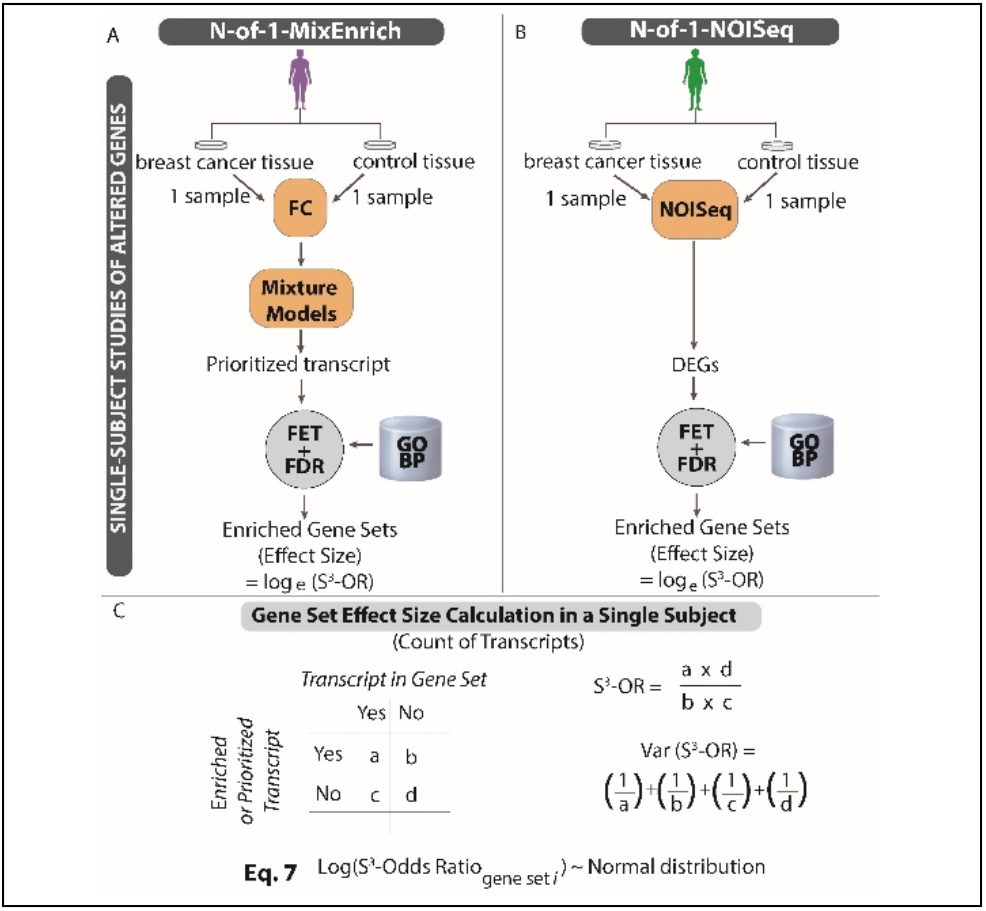
Overview of two single-subject study methods conducted from one sample per condition without replicate generating effect sizes and variance for each gene set. We apply single-subject studies to each subject to identify either prioritized transcripts (**Panel A**) or DEGs (**Panel B**) between paired tumor-normal samples. We identify patient specific enriched gene sets and associated effect sizes in the form of natural log odds ratios through a FET (**Panel C**). Each effect size is approximately normally distributed with known variance and mean, simplifying subsequent analyses between cohorts. The gene set-level variance enables the extraction of more information from each individual subject than typical variance estimators that work across subjects and thereby leads to increased statistical power. The N-of-1-MixEnrich method was previously described and validated (Berghout, et al., 2018; Li, et al., 2017; Zaim, et al., 2018). NOISeq is also considered as an alternative meriting evaluation because of its performance in prior single-subject studies evaluations (Zaim, et al., 2019).

We also applied NOISeq to each of the tumor-normal pairs (Tarazona, et al., 2015) as shown in **Panel B** of **Figure 2**. For these applications of NOISeq with no replicates, the “*pnr*” and “*ν*” parameters were set to 0.0002 and 0.00002 to prevent the method from producing any errors related to setting the size of the inherent multinomial distributions to an integer too large for R to handle. The criteria for identifying genes as differentially expressed for NOISeq were the same as those used for N-of-1-MixEnrich. As shown in **Panel C** of **Figure 2** (next page), we subsequently used this information to construct contingency tables and calculate the natural log odds ratio for *Inter-N-of-1*. This process generated two different applications of *Inter-N-of-1*, N-of-1-MixEnrich and NOISeq, to conduct the single-subject analyses preceding the cohort comparison.

#### Comparing Enriched *Gene Sets* across Distinct Cohorts

We first combined the data within two distinct cohorts into single statistics whose null reference distributions were at least approximately normal. These within-cohort statistics were contrasted via scaled subtraction in a manner reminiscent of the two-sample t-test to establish the difference in gene set enrichment between the two cohorts. Let *gs* ∈ {1,…, *N*] index the specific ***g**ene **s**et* being studied where *N* is the total number of gene sets, *k_j_* indexes a specific subject in cohort *K* composed of *S_K_* individuals with subjects numbered *j* ∈ {1,…, *S_K_*}, and *K* ∈ {*A, B*} indexes a specific cohort. Let ***Δ*** signify quantities relating to the difference between the two cohorts.

The *Inter-N-of-1* analytics for combining information within a cohort considers the abstract contingency table shown as **Table 3** where the cell counts are representative for the gene set indexed by *gs* and the subject indexed by *k_j_*.

**Table 3:**
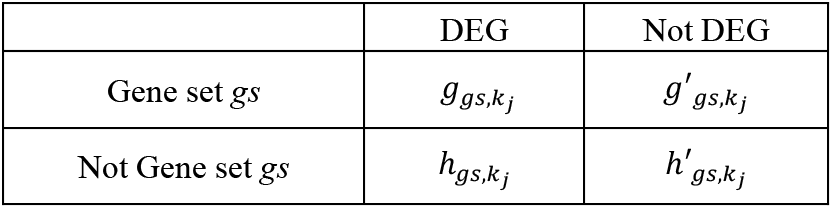
Notation for 2×2 Contingency Table Cross-classifying DEG Status with Gene Set Status

We obtain DEGs from the application of a chosen single-subject analysis method (either N-of-1-MixEnrich or N-of-1-NOISeq) for a specific gene set *gs* in individual *k_j_* of cohort *K* to fill out the contingency table with counts in the format shown in **Table 3**. We apply a continuity correction by adding 0.5 to each of the cells in the contingency table to provide a small-sample adjustment in the odds ratio (Agresti and Kateri, 2011). The natural log S^3^ OR, denoted as *Q_gs,k_j__*, **Equation (5)**, is approximately normally distributed with variance var (*Q_gs,k_j__*) given in **Equation (6)** (Woolf, 1955).

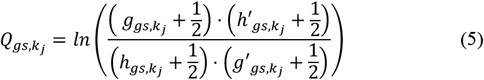

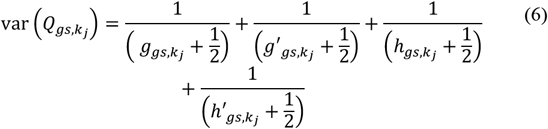

We average the *Q_gs,k_j__* values within their respective cohorts to obtain the average ln ORs

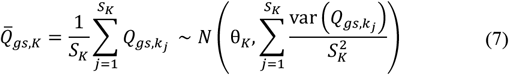

When the null hypothesis *H*_0_:*θ_A_* = *E*[*ln*(OR_*A*_)] = *E*[*ln*(OR_*B*_)] = θ_*B*_ is true then

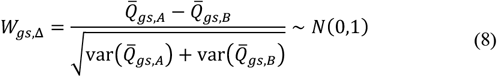

at least approximately. The corresponding two-sided p-value for gene set *gs* is

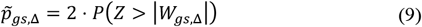

where Z represents a standard normal random variable. An FDR adjustment via the Benjamini-Hochberg method (Benjamini and Hochberg, 1995) is then applied to the 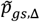 across all the GO terms tested in the particular application. To ensure that the method positively identifies gene sets that are enriched in at least one of the cohorts, we set all FDR adjusted p-values to 1.0 if both cohort means of the log odds ratios are negative. This step ensures interpretable results since impoverished GO terms with significantly fewer-than-expected DEGs are not well understood from a biological context.

#### 2.3 Description of the Generalized Linear Models and application of *Inter-N-of-1* methods for small cohort comparison and their evaluation in the Breast Cancer Data

##### Generalized Linear Model (GLM) Designs

For the cohort analyses, we applied a generalized linear model as implemented in *limma (Smyth, et al., 2005)*. Preceding application of the generalized linear model, we performed trimmed mean of M values (TMM) normalization (Robinson and Oshlack, 2010) on the data pre-processed for cohort analysis. We applied the voom normalization (Law, et al., 2014) via the *limma* function *voom-withQualityWeights* in R.

We used the three different designs described in **Table 4** for these generalized linear model-based analyses, which were called the simple design, the interaction design, and GLM+EGS respectively. We blocked by subject for each of these GLM designs, and all FDR adjustments of p-values were done using the Benjamini-Hochberg False Discovery Rate (FDR) method (Benjamini and Hochberg, 1995).

**Table 4.** Three experimental designs used for the generalized linear models. In the analysis of subsets of the TCGA Breast Cancer data, genes were declared differentially expressed if their abs(log_2_FC) > log_2_(1.2) and their FDR-adjusted p-value < 0.05. Within the simulation, genes were declared differentially expressed if their FDR-adjusted p-values < 0.05.

| Name | Level | What is compared | Results |
| --- | --- | --- | --- |
| Simple | Transcript | TP53_Tumoral – PIK3CA_Tumoral | **Fig. 3 Panel A** |
| Interaction | Transcript | (TP53_Tumoral – TP53_Normal) – (PIK3CA_Tumoral – PIK3CA_Normal) | **Fig. 3 Panel B** |
| GLM+EGS | Gene set | 1) Find DEGs using Interaction Contrast<br>2) Enrichment via FET | **Fig. 4 - 5** |

##### Reference standard construction of enriched pathways using *edgeR* Generalized Linear Model followed by Gene Set enrichment

After pre-processing for cohort analyses, we applied generalized linear models as implemented in the R software package *edgeR* (Robinson, et al., 2010) at FDR< 5% to the entire TCGA breast cancer dataset to construct three reference standards corresponding to the three designs discussed in **Table 4**. Each reference standard evaluated the analyses of the TCGA breast cancer cohorts (TP53 vs PIK3CA) and used the same filter thresholds for classifying transcripts as differentially expressed. In the GLM followed by enrichment of gene set (GLM+EGS), the prioritized interacting transcripts are followed by a FET at FDR<5%.

##### Subsampling of the TCGA Breast Cancer Cohort and application of GLM and *Inter-N-of-1* methods

For each of the values *S_A_* = *S_B_* = *S* ∈ {2,3,4,5, 7,8,9} we ran 100 subsamples of the total cohorts where we randomly selected without replacement S subjects with *TP53* and S subjects with *PIK3CA*, without requiring non-redundancy of the random samplings. We applied the GLM+EGS method and the N-of-1-MixEnrich and NOISeq versions of the *Inter-N-of-1* method to each of the selected cohorts (TP53 vs PIK3CA). For each of the three methods, FDR<5% adjustment of the p-values was done with respect to all 5,179 GO terms tested.

For random subsamples of size *S_A_* = *S_B_* = *S* ∈ {2,3,4,… 19} of subjects, we applied the two transcript-level analyses using generalized linear models as implemented in *limma*. The performance of these transcript-level applications of *limma* were assessed and illustrated in **Figure 3** to demonstrate the necessity and benefit of transforming from transcript-level to gene set-level analyses.

**Fig. 3.**
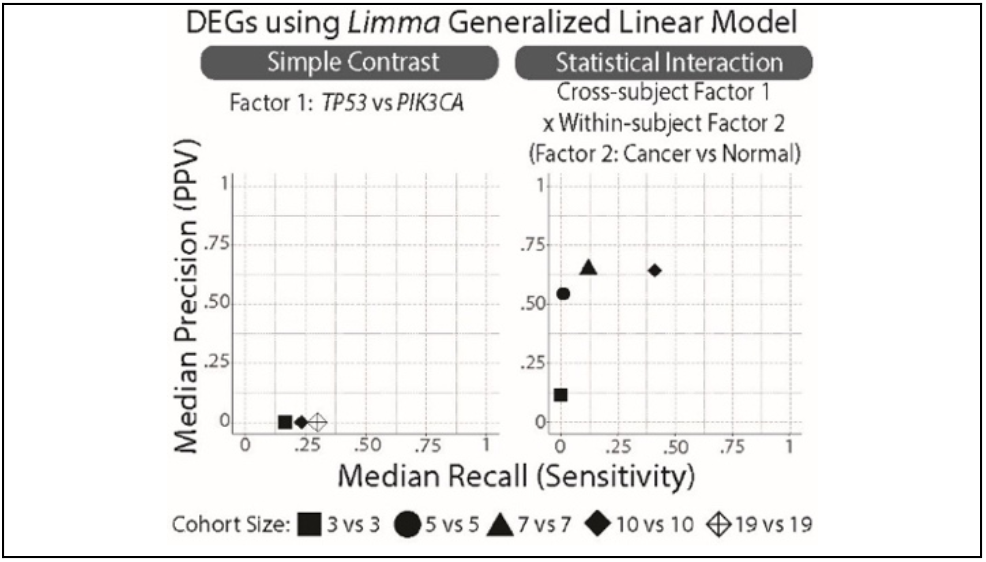
At the transcript level, limited accuracies of Generalized Linear Models for calculating conventional simple contrast or interactions in small heterogenic breast cancer cohorts. While GLMs can deliver DEGs in small cohorts for isogenic cellular and animal models, we recapitulate in the TCGA datasets that small human cohorts are underpowered statistically. We calculated the precision and recall scores associated with each of the 100 random sub-samplings of cohort sizes 2vs2, 3vs3, …, to 19vs19 for *TP53* vs *PIK3CA* and report median accuracie*s*. The left panel used a simple linear contrast of the tumor levels on the molecular subtypes. The right panel used a linear contrast corresponding to the interaction between the molecular subtypes (TP53 vs PIK3CA) and tumor status (Breast cancer vs normal breast). Discoveries were performed with *limma* while the reference standard was constructed with *edgeR*.

##### Accuracy measures within TCGA breast cancer dataset

For each method, we calculated the precision and recall using the following functions. When a method produced no positive predictions for the gene sets, we assigned values of zero to the precision and recall of the given method. Otherwise, we calculated the precision and recall using Powers’ calculations with adjustments of adding 0.5 to numerators and 1.0 to denominators to avoid divisions by zero (Powers, 2020). In addition, we have previously published extensions to conventional accuracy scores that we termed “similarity Venn Diagrams” and “Similarity Contingency Tables” (Gardeux, et al., 2015). In these approaches, identical as well as highly similar GO-BP terms between the prediction set and the reference standard account for true positive results. We calculated the precision and recall of the gene set level analyses using Information Theoretic Similarity (ITS) (Tao, et al., 2007). For precision, we included in the intersection those predicted GO-BPs which had an ITS similarity of 0.70 or higher with any of the GO terms in the reference standards, while the denominator remained as all predicted GO-BPs. Similarly, for recall we included in the intersection the reference standard GO-BPs which had an ITS similarity score of 0.70 or higher with any of the predicted GO terms, while the denominator remained as the total positive reference standard GO-BP terms. Of note, we previously reported that this ITS>0.70 similarity criteria is highly conservative since ~0.0056 pairs of GO-BP terms are similar at ITS>0.7 (58,577 pairs among 10,458,756 non-identical combinations of GO-BPs) (Gardeux, et al., 2015).

#### 2.4 Simulation of small cohort comparisons to compare GLMs to *Inter-N-of-1* methods

##### Data generation for Simulation

The overall scheme for the simulation began by constructing two cohorts of paired tumor-normal RNA-seq expression profiles. We calculated simulation parameters to most realistically create these expression values as described below (**Table 5**). To calculate statistical interactions between two factors, we had to design two cohorts of subjects and each subject with two sample conditions. We sought to recreate the TCGA Breast cancer conditions with these parameters, using the observed median values in the TCGA dataset as the medians of the simulation parameters and varying the parameters around said medians. The TCGA dataset did not comprise repeated samples in the same condition, and thus we utilized the unstimulated MCF7 cell lines with seven replicates to estimate the variation expected between two paired normal tissues. In our previous pathway expression studies ((Yang, et al., 2012) and data not shown) where we compared two cohorts, about two-thirds of the observed responsive gene set patterns - as shown in **Figure 2** - consisted of a gene set responsive in one subject cohort and unresponsive in the other cohort.

**Table 5.** Simulation Parameter Values. Only the balanced cohort size and the proportion of subjects with coordinated DEGs were varied. All other parameters were held constant. 30 datasets were generated for each parameter configuration leading to a total of 540 datasets.

| Parameters | How Estimated | Values |
| --- | --- | --- |
| Control Samples | Randomly sample without replacement from TCGA breast cancer normal samples | NA |
| log<sub>2</sub>FC distribution of non-differentially expressed genes | 1) Calculate log<sub>2</sub>FCs of randomly paired MCF7 unstimulated breast cancer samples<br>2) Split log<sub>2</sub>FCs into deciles by baseline expression<br>a) All deciles containing 0 are combined into one category<br>3) Sample with replacement from decile containing transcript name in first random pair | NA |
| Gamma parameters of log<sub>2</sub>FCs of DEGs | 1) Run N-of-1-MixEnrich (Fig.2) on within-subject tumor-normal pairs in TP53 and PIK3CA cohorts to identify DEGs<br>2) MLEs for gamma parameters fit to absolute log<sub>2</sub>FCs of DEGs<br>a) Used *egamma* function in *EnvStats R* package (Millard, et al., 2020) | Scale parameter = 6.06<br>Shape parameter = 0.55 |
| Proportion of DEGs in enriched GO-BPs | 1) Split enriched GO terms from *edgeR* reference standard into deciles based on size<br>2) Calculated DEGs median proportion for deciles containing GO-BPs (size: 47, 200) | (GO size 200): 0.10<br>(GO size 40): 0.19 |
| Proportion of Subjects with coordinated DEGs | 1) Split log<sub>2</sub>FCs of DEGs within *edgeR* reference standard into categories<br>a) >1.3, b) < -1.3, or c) neither<br>2) Assign the maximum proportion of subjects per categories (a) or (b) for each transcript<br>3) Find the median proportion of subjects across all transcripts | 0.25, 0.48, 0.75 |
| Balanced Cohort Size | NA | 2, 3, 7, 10, 20, 30 |
| GO-BP terms | 1) Enriched: GO:0002221 (200 genes)<br>2) Enriched: GO:0000096 (47 genes)<br>3) Control: GO:0006733 (196 genes)<br>4) Control: GO:0090184 (41 genes) | NA |

These paired tumor-normal samples represented within-subject samples were constructed to have a proportion of the transcripts with altered expression between the tumor and normal states. Through the use of randomly sampling without replacement, we generated the normal tissue samples for these pairs after filtering out all genes in the 112 TCGA breast cancer normal tissues, which were not present within the MCF7 breast cancer dataset (leaving 17,414 genes).

For each sampled normal breast tissue sample, we generated transcript expression for a paired breast cancer sample of that subject rather than sampling the corresponding breast cancer sample from the TCGA data. To produce a paired tumor expression value for a non-differentially expressed gene, we first followed the steps outlined in **Table 5** to randomly generate empirical log_2_ Fold Changes (log_2_FC) and then we set the gene’s expression as the product of the gene’s paired normal expression and 2 raised to the exponent of the log_2_FC value. To generate the expression value for an altered transcript in a tumor sample, we randomly sampled a log_2_FC from a gamma distribution with parameters described in **Table 5** and set said gene’s expression to the product of the gene’s normal expression and 2 raised to the exponent of the log_2_FC value. We generated only positive log_2_FCs for the DEGs to improve the *GLM’s* ability to detect them as differentially expressed cross subjects. We specified a gamma distribution for these positive log_2_FCs since all the absolute-valued log_2_FC distributions we examined possessed significant right-skew.

We chose to evaluate the methods using the 4 GO terms described in **Table 5**. In simulation cohort A, 2 of these GO-BPs would be seeded with altered transcripts, thus enriched, and 2 would serve as controls. In cohort B, none of the 4 GO terms were enriched, thereby setting up an interaction effect between the within-subject and between-subject factors. Within the two enriched GO terms in cohort A, we randomly selected the proportions of genes specified in **Table 5** to have altered expression. We used Bernoulli random variables with probabilities of success outlined in **Table 5** to designate subjects within cohort A, which would share all their randomly selected DEGs. The remaining subjects within cohort A had all their DEGs randomly vary across subjects. It was hypothesized that the percentage of subjects with shared altered transcripts would strongly influence the performance of the GLM+EGS method since *limma* assumes the presence of coordination of gene expression across subjects. Thus, we varied the expected proportion of subjects with shared DEGs within cohort A (0.25, 0.48, 0.75) along with the sizes of the two cohorts (2, 3, 7, 10, 20, 30) while holding all other parameters constant. We consequently generated 30 datasets for each parameter combination leading to a total of 540 datasets for our downstream simulations.

##### Data preprocessing within Simulation

(A) For the generalized linear model analyses, we preprocessed the simulated data by removing all genes with mean expression values less than 30 across all the simulated transcripts and subsequently added 1 to each of the expression counts. (B) For the single-subject analyses, we applied a two-stage pre-processing method in which we (i) removed all the transcripts with mean expression less than 30 within each sample-pair and (ii) found the union across all pairs of genes remaining and eliminated any genes not contained within. The remaining genes for the single-subject analyses then had 1 added to their expression counts to eliminate any remaining zeroes.

##### Application of Methods to Simulated Data

The GLM+EGS and the two versions of the *Inter-N-of-1* method were applied to each of the generated datasets as described previously. The Benjamini-Hochberg False Discovery Rate (FDR) (Benjamini and Hochberg, 1995) adjustments of the p-values generated for each technique were performed with respect to only the 4 selected GO terms that were tested for each combination of dataset and method. GOBPs were declared positive for a method if their associated FDR adjusted p-values for said method were below 0.05.

##### Accuracy measures within the Simulation

To estimate the overall performance of each method within the simulation, we calculated the number of true positives, true negatives, false positives, and false negatives occurring within the 2 enriched and 2 control GO terms across all 30 resampling of each combination of parameters. When any of the methods made no positive predictions for the gene sets, we artificially assigned values of 0 to the precision and recall of the given method. Otherwise, we calculated the precision and recall through the use of their traditional formulae (Powers, 2020). 30 accuracy scores are thus available for each combination of parameters for each GO term size (40 and 200).

### 3 Results

We showed that using a two-step process, where we first enrich the signal-to-noise ratio by applying S^3^-analyses to paired data in single-subjects before combining across subjects, can capture stable signal and yield results comparable to those in the reference standard, even as cohort size decreases. By contrast, traditional techniques for identification of gene set-level biomechanisms that differentiate between two cohorts rapidly lose power and yield unreliable results as the sample size decreases below 5 subjects per cohort.

The transcriptomic analyses of TCGA data in **Figure 3** recapitulates that small human cohorts are particularly difficult to analyze using GLMs due to their heterogenic conditions and lack of controlled environment. Thus, small human cohorts present a stark contrast to isogenic controlled experiment cell lines or animal models where the high signal to noise ratio makes transcriptomic analyses possible for very small sample sizes. These unsurprising results provide the justification for the development of the proposed GLM+EGS and *Inter-N-of-1* methods conducted at the gene set level. They also attest to the intrinsic lack of signal within the TCGA breast cancer data for such transcriptomic analyses.

The performance results for subsets of the TCGA breast cancer data shown in **Figure 4** establish that the two versions of the proposed *Inter-N-of-1* method degrade more gracefully in performance with decreasing cohort size than traditional generalized linear model-based methods, thereby allowing them to outperform for smaller cohort sizes. **Figure 4** shows that the niche where the *Inter-N-of-1* methods outperform in terms of median precision and recall extends to all cohort sizes below 7vs7, with the GLM+EGS method achieving higher median performance scores for 9vs9 and above. The sizes of the crosses suggest a further boon for the developed methods beyond this better ‘on average’ performance. The *Inter-N-of-1* methods tend to have very small tight crosses suggesting low variation in performance and greater consistency. The GLM+EGS method on the other hand possesses very large crosses until cohort size 9vs9, suggesting wild swings in performance across the different subsets evaluated. In addition, even the gene set-level GLM+EGS method outperforms transcript-level GLM analyses (**Fig. 3 vs Fig. 4**). **Figure 4** also establishes that the N-of-1-MixEnrich version of the *Inter-N-of-1* method outperforms the NOISeq version in terms of consistency and median precision and recall. Although these differences remain small for larger cohort sizes of 7vs7 and above, they increase gradually with decreasing cohort sizes.

**Fig. 4.**
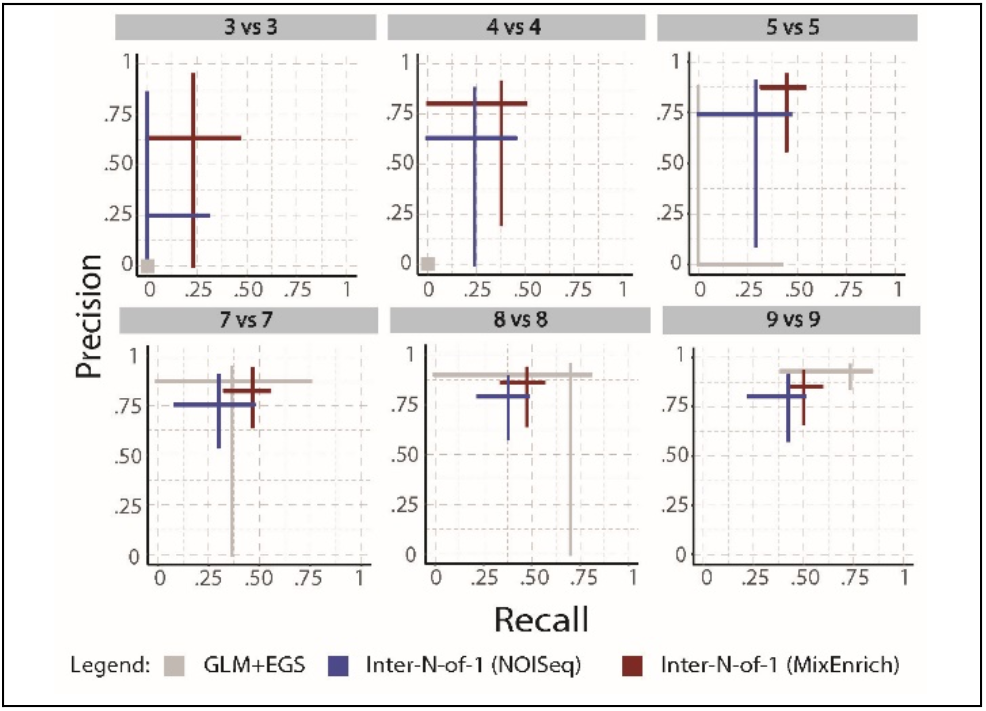
At the gene set–level, two *Inter-N-of-N* methods outperform a GLM followed by enrichment in small heterogenic human cohorts. While *Inter-N-of-1* methods (*Inter-N-of-1* (NOISeq) and *Inter-N-of-1* (MixEnrich)) outperform the GLM followed by enrichment in gene sets for sample sizes of 7vs7 and smaller, the GLM+EGS shows better accuracy at sample sizes 9vs9 and above. Of note, GLM+EGS shows large variations in performance measures within the samples of size 8vs8 suggesting that despite its improved median accuracy it remains unreliable at that level. In all cases, the discovery of differentially responsive gene sets (*Inter-N-of-1* methods) or enriched gene sets (GLM+EGS) substantially outperform the accuracies of transcript-level analyses shown in **Fig. 3**. While the *Inter-N-of-1* and GLM+EGS methods identify related signals, the reference standard designed by a distinct GLM+EGS approach favors the accuracies of the latter. In addition, *Inter-N-of-1* methods can assess the effect size of responsive gene sets in each subject, which can be illustrated as box plots of gene set response. In contrast. GLM+EGS methods are limited to a single description of over-representation calculated on interacting transcripts of the entire study. We calculated the precision and recall scores associated with each of the 100 random subsampling of cohort sizes 2vs2, 3vs3, 4vs4, 5vs5, 7vs7, 8vs8, 9vs9 for *TP53* and *PIK3CA* subjects with the GLM+EGS and *Inter-N-of-1* methods: (i) *Inter-N-of-1* (*NOISeq*), and (ii) *Inter-N-of-1* (*MixEnrich*). The arms extend from the lower quartile to the upper quartile of the respective performance measure, and the two arms cross at the median for the precision and recall for that technique at the indicated cohort size.

The simulations indicates that the proposed *Inter-N-of-1* methods outperform GLM+EGS for small sample sizes within parameters derived from cancer datasets and extended to. investigate other conditions. **Fig. 5** shows that the two *Inter-N-of-1* methods are unaffected by changes in the expected proportion of subjects within cohorts with shared DEGs since their performance scores typically oscillate randomly around a fixed point given a fixed cohort size. These fixed points come closer to the perfect score of 1.0 precision and 1.0 recall with increasing cohort size, suggesting that mainly the cohort size affects the *Inter-N-of-1* method. The N-of-1-MixEnrich version of the *Inter-N-of-1* method generally performs the best out of all three methods, with its precision always staying 90% or higher and its recall staying 75% or above for all parameter configurations. The NOISeq version of the *Inter-N-of-1* method suffers from a higher rate of false negatives for the two smallest tested cohort sizes of 2 and 3 and so displays significantly less recall than the N-of-1-MixEnrich version of the *Inter-N-of-1* method, although it does display similar levels of precision. Thus, this simulation also unveils the reason for which *Inter-N-of-1* (NOISeq) did not perform as well. Both cohort size and the expected proportion of subjects within groups with coordinated DEGs affect the performance of the GLM+EGS method. Increasing either of these parameters significantly improves the performance of the GLM+EGS method, with the single exception of the 2vs2 cohort size where GLM+EGS produces 0 precision and recall for all specifications of the proportion of subjects within group with coordinated DEGs. At the anti-conservative levels for these parameters, the GLM+EGS method matches the performance of the two versions of the *Inter-N-of-1* method. However, decreasing either parameter quickly leads the GLM+EGS method to underperform. For cohort sizes of 10vs10 and lower, the GLM+EGS method fails to match the performance of the two versions of the *Inter-N-of-1* method and so supports the superiority of *Inter-N-of-1* in such small sample sizes for breast cancer-like data.

**Figure 5.**
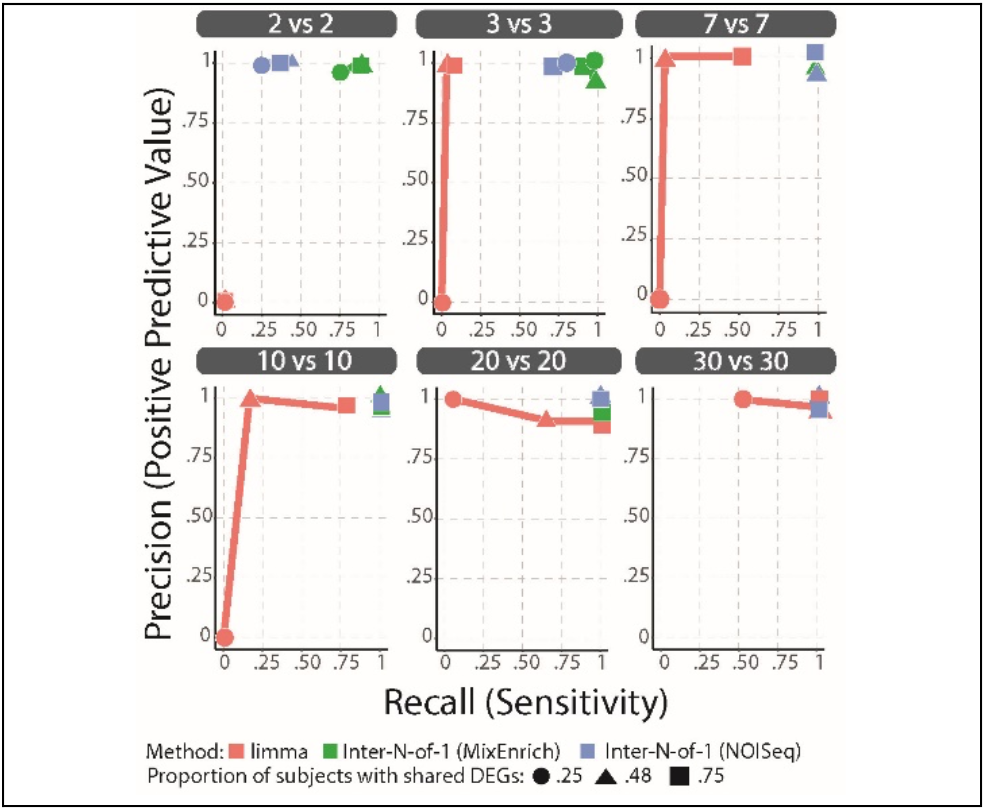
Comparison of accuracy of GLM+EGS and *Inter-N-of-1* methods within the simulation. We generated subject tumor-normal pairs for a variety of cohort sizes (2vs2, 3vs3, 7vs7, 10vs10, 20vs20, 30vs30) and expected proportion of subjects with shared DEGs in cohort A (0.25, 0.48, 0.75). We simulated 30 datasets for each parameter configuration and applied the proposed developed *Inter-N-of-1 methods* and GLM+EGS method to each. We calculated the total number of true positives, false positives, false negatives, and true negatives across all iterations and used them to calculate the precision and recall for each combination of method, parameter configuration, and GO term size. Separate graphs are made for each parameter configuration and plot the resulting precision and recall measures for each method for the gene sets of size 40. The results for gene sets of size 200 were very similar to the above results and so were excluded. The *N-of-1-MixEnrich* version of the *Inter-N-of-1* method performs excellently and achieves near perfect scores for cohort sizes above 2. The NOISeq version of the *Inter-N-of-1* method often fails to identify positive signal for cohort sizes of 3 or smaller, but otherwise achieves performance scores near those of the *N-of-1-MixEnrich* version of the *Inter-N-of-1* method. The two versions of *Inter-N-of-1* appear to be unaffected by changes in the expected proportion of subjects with shared DEGs since their performance scores within each graph oscillate around the same general area and show no overall trend. The GLM+EGS method often struggles to identify positive signal for smaller cohort sizes, although increasing the expected proportion of subjects within cohorts with coordinated DEGs improves the recall of the method and decreases the minimum sample size needed for it to perform near perfectly. The GLM+EGS method always shows excellent precision and control of the overall FDR for all except the cohort sizes of 2.

## 4 Discussion

As stated in the introduction, empirical evidence suggests the existence of a methodological gap when comparing transcriptomic differences in biomechanisms within very small human cohorts due to variations of heterogenicity, uncontrolled biology (age, gender, etc.), and diversity of environmental factors (nutrition, sleep, etc.).As expected, state of the art generalized linear models decline in performance with sample sizes less than 5 (Soneson and Delorenzi, 2013). Smaller datasets require variances to be as low as those observed between technical replicates or with the isogenic conditions of cellular and animal models. Yet, even in such isogenic conditions, two studies have recommended at least 6 biological replicates for applying generalized linear models (Liu, et al., 2014; Schurch, et al., 2016). Examining two-factor interactions in transcriptomes (Cohorts × tumor status) further inflates the required sample size by a factor of 4 (Brookes, et al., 2004; Fleiss, 2004; Leon and Heo, 2009). Traditional cohort-based methods impose sample size requirements which simply cannot be met within the framework imposed by rare diseases, prompting the need to develop new methods.

On the other hand, we and others have shown it is possible to obtain statistical significance of gene set-level effect size measures from single samples without replicates taken in two conditions, namely single-subject studies (S^3^) (Li, et al., 2017; Li, et al., 2017; Schissler, et al., 2015; Vitali, et al., 2017). We have shown evidence from breast cancer studies and simulations that the S^3^-anchored *Inter-N-of-1* addresses this methodological gap. Their slow decay in performance when contrasted with the abrupt decay of GLM+EGS establishes the superiority of these methods for sample sizes of *S_A_* = *S_B_* ∈ {2,3,4,5, 6} when applied to our TCGA breast cancer dataset. Comparison of the median precision and recall of the three considered techniques shows that on average our methods exhibit greater power and importantly less variable performance than GLM+EGS at these low cohort sizes. Furthermore, our simulation study confirmed that both versions of the *Inter-N-of-1* provide substantially improved recall over the GLM+EGS method at small cohort sizes while still maintaining equivalent levels of precision. The simulation results also establish that the expected proportion of subjects with coordinated DEGs within cohorts plays a critical role in determining the range of cohort sizes in which the developed methods outperform traditional generalized linear model-based techniques. In datasets where the proportion of subjects within cohorts sharing their DEGs is lower than 48%, the *Inter-N-of-1* methods continue to out perform the GLM+EGS method for cohort sizes larger than 20.

Several limitations were observed. (1) This study focuses on parameters related to cancers, where there are substantial differences between normal paired tissue to cancer tissue. While single-subject studies have been shown to be effective in viral response (Gardeux, et al., 2017; Gardeux, et al., 2015) or response to therapy (Li, et al., 2017), it remains to be demonstrated that the downstream *Inter-N-of-1* methods can outperform transcript-level methods in those biological conditions. (2) The simulation does present some inconsistencies with observations made within the TCGA breast cancer subsets. This can probably be explained by the fact that the breast cancer analyses used a reference standard that favored GLM+EGS over *Inter-N-of-1* methods by design. (3) We explored only one type of difference within gene set response between cohorts in the simulations: a cohort responsive vs unresponsive. We are thus undertaking the complementary analysis to compare the more general paradigm of gene sets more responsive in one cohort than in the other. (4) Finally, although the developed methods allow for a more accurate testing of interactions in datasets with small sample sizes, the importance of balancing confounders between the two cohorts should not be overstated. The small samples used within these analyses prevent randomization from balancing key covariates and confounders between cohorts. Future studies could model unbalanced covariates through data or knowledge fusion with external datasets. (5) Transcript independence assumptions in the calculation of the single-subject odds ratio and its variance (*Inter-N-of-1* methods) may be transgressed. However, many such assumptions are routinely overlooked in related analyses, such as BH-FDR (Benjamini and Hochberg, 1995) with similar limitations later rectified as the BY-FDR (Benjamini and Yekutieli, 2001). When viewed under that perspective, computational biology may progress by first proving new models and then addressing their biases in subsequent studies. (6) Other unbiased approaches to generating gene sets could have been utilized (e.g., co-expression network from independent datasets, protein interaction networks, etc.). (7) Of note, few datasets are available with two measures in different conditions per subject and more than one clinical cohort of subjects. Similar to physics where experimentalist and theory influence one another, our work presents improvements on solving an experimental design that is infrequently used and merits more consideration for increasing the signal-to-noise ratio in the study of rare and infrequent diseases. (8) Prospective biologic validation of results is also required in future studies as we have done with single-subject studies in the past (Gardeux, et al., 2014).

Another consideration concerns that how GLM+EGS and *Inter-N-of-1* evaluate different phenomena. The GLM+EGS method primarily discovers GO terms enriched for transcripts – primarily require the coordination of signals at the transcript-level before the enrichment across subjects belonging to similar classness. The *Inter-N-of-1*, on the other hand, assesses whether the proportion of responsive transcripts within a given GO term measured in each subject significantly differs across cohorts at the gene set-level. In other words, in the *Inter-N-of-1*, the transcripts contribution to the gene set signal may be different between subjects, while in the GLM_EGS methods a transcript-level coordination is required. The *Inter-N-of-1* favors clinical applications where gene set mechanisms are causal to the disease. Cancer is one such condition where numerous genetic and transcriptomic root causes may differ between subjects and yet converge to comparable cellular and clinical phenotypes.

In conclusion, the proposed S^3^-anchored Effect Size-methods demonstrate the utility of within-subject paired sample designs for better controlling within-patient background genetic variation and thereby identifying clearer signal with small numbers of subjects. These approaches first simplify the heterogenicity between subjects with better controlled single-subject studies reminiscent of experimental isogenic models (e.g., cell lines or mice models). These results motivate further studies of new experimental designs, where paired within-subject samples allow analyses of datasets previously considered too small. The new design not only presents opportunities in terms of performance within small subject cohorts, but also in terms of utility. The use of single-subject methods within the *Inter-N-of-1* creates an avenue for examining subject variability within cohorts. By examining the single-subject results one can directly see the degree of concordance and discordance amongst subjects and answer questions pertaining to whether specific subjects possess the overall observed signal. Thus, the *Inter-N-of-1* presented here represents not just a new method that performs better within small sample sizes, but also an example for how to borrow knowledge from gene sets for more powerful measures of dispersion in a single subject to conduct studies of rare or infrequent diseases and analyses on patient variability within and across cohorts. In addition, precision therapies designed for increasingly substratified common disorders can benefit from the proposed methods. The strategies and methods presented here open a new frontier that may greatly enrich our understanding of the genetic foundations of rare diseases.

## Acknowledgements

We acknowledge Branden Lau for performing alignment of the MCF7 SRA files.

## References

Agresti, A. and Kateri, M. Categorical data analysis. Springer Berlin Heidelberg; 2011.

Andre, F., et al. Alpelisib for PIK3CA-Mutated, Hormone Receptor-Positive Advanced Breast Cancer. N Engl J Med 2019;380(20):1929–1940.

Ashburner, M., et al. Gene ontology: tool for the unification of biology. Nature Genetics 2000;25(1):25.

Balli, M., et al. Autologous micrograft accelerates endogenous wound healing response through ERK-induced cell migration. Cell Death & Differentiation 2019:1–19.

Benjamini, Y. and Hochberg, Y. Controlling the False Discovery Rate - a Practical and Powerful Approach to Multiple Testing. JR StatSocB 1995;57(1):289–300.

Benjamini, Y. and Yekutieli, D. The control of the false discovery rate in multiple testing under dependency. Annals of statistics 2001:1165–1188.

Berghout, J., et al. Single subject transcriptome analysis to identify functionally signed gene set or pathway activity. In, PSB. World Scientific; 2018. p. 400–411.

Berghout, J., et al. Single subject transcriptome analysis to identify functionally signed gene set or pathway activity. Pac Symp Biocomput 2018;23:400–411.

Brookes, S.T., et al. Subgroup analyses in randomized trials: risks of subgroup-specific analyses;: power and sample size for the interaction test. Journal of Clinical Epidemiology 2004;57(3):229–236.

Cancer Genome Atlas, N. Comprehensive molecular portraits of human breast tumours. Nature 2012;490(7418):61–70.

Ciriello, G., et al. Comprehensive Molecular Portraits of Invasive Lobular Breast Cancer. Cell 2015;163(2):506–519.

Edgar, R., Domrachev, M. and Lash, A.E. Gene Expression Omnibus: NCBI gene expression and hybridization array data repository. Nucleic acids research 2002;30(1):207–210.

Elliott, E.J. and Zurynski, Y.A. Rare diseases are a’common’problem for clinicians. Australian family physician 2015;44(9):630.

Fisher, R.A. The logic of inductive inference. Journal of the Royal Statistical Society 1935;98(1):39–82.

Fleiss, J. The design and analysis of clinical experiments. 1986. New York, John Wiley & Sons 2004.

Gardeux, V., et al. Concordance of deregulated mechanisms unveiled in underpowered experiments: PTBP1 knockdown case study. BMC medical genomics 2014;7(1):1–13.

Gardeux, V., et al. Concordance of deregulated mechanisms unveiled in underpowered experiments: PTBP1 knockdown case study. BMC Med Genomics 2014;7 Suppl 1(S1):S1.

Gardeux, V., et al. A genome-by-environment interaction classifier for precision medicine: personal transcriptome response to rhinovirus identifies children prone to asthma exacerbations. Journal of the American Medical Informatics Association 2017;24(6):1116–1126.

Gardeux, V., et al. Towards a PBMC “virogram assay” for precision medicine: Concordance between ex vivo and in vivo viral infection transcriptomes. Journal of biomedical informatics 2015;55:94–103.

Griggs, R.C., et al. Clinical research for rare disease: opportunities, challenges, and solutions. Mol Genet Metab 2009;96(1):20–26.

Grossman, R.L., et al. Toward a Shared Vision for Cancer Genomic Data. N Engl J Med 2016;375(12):1109–1112.

Guillem, P., et al. Rare diseases in disabled children: an epidemiological survey. Arch Dis Child 2008;93(2):115–118.

Kim, J.Y., et al. Clinical implications of genomic profiles in metastatic breast cancer with a focus on TP53 and PIK3CA, the most frequently mutated genes. Oncotarget 2017;8(17):27997–28007.

Law, C.W., et al. voom: Precision weights unlock linear model analysis tools for RNA-seq read counts. Genome biology 2014;15(2):R29.

Leon, A.C. and Heo, M. Sample sizes required to detect interactions between two binary fixed-effects in a mixed-effects linear regression model. Computational statistics & data analysis 2009;53(3):603–608.

Li, Q., et al. N-of-1-pathways MixEnrich: advancing precision medicine via single-subject analysis in discovering dynamic changes of transcriptomes. BMC Med Genomics 2017;10(Suppl 1):27.

Li, Q., et al. kMEn: Analyzing noisy and bidirectional transcriptional pathway responses in single subjects. J Biomed Inform 2017;66:32–41.

Liu, Y., Zhou, J. and White, K.P. RNA-seq differential expression studies: more sequence or more replication? Bioinformatics 2014;30(3):301–304.

Millard, S.P., Kowarik, A. and Kowarik, M.A. Package ‘EnvStats’. 2020.

Ozturk, K., et al. The Emerging Potential for Network Analysis to Inform Precision Cancer Medicine. J Mol Biol 2018;430(18 Pt A):2875–2899.

Powers, D.M. Evaluation: from precision, recall and F-measure to ROC, informedness, markedness and correlation. arXiv preprint arXiv:2010.16061 2020.

Robinson, M.D., McCarthy, D.J. and Smyth, G.K. edgeR: a Bioconductor package for differential expression analysis of digital gene expression data. Bioinformatics 2010;26(1):139–140.

Robinson, M.D. and Oshlack, A. A scaling normalization method for differential expression analysis of RNA-seq data. Genome Biology 2010;11(3):R25.

Schissler, A.G., et al. Dynamic changes of RNA-sequencing expression for precision medicine: N-of-1-pathways Mahalanobis distance within pathways of single subjects predicts breast cancer survival. Bioinformatics 2015;31(12):293–302.

Schissler, A.G., et al. Analysis of aggregated cell–cell statistical distances within pathways unveils therapeutic-resistance mechanisms in circulating tumor cells. Bioinformatics 2016;32(12):i80–i89.

Schissler, A.G., Piegorsch, W.W. and Lussier, Y.A. Testing for differentially expressed genetic pathways with single-subject N-of-1 data in the presence of inter-gene correlation. Stat Methods Med Res 2018;27(12):3797–3813.

Schurch, N.J., et al. How many biological replicates are needed in an RNA-seq experiment and which differential expression tool should you use? Rna 2016;22(6):839–851.

Smyth, G.K., et al. LIMMA: linear models for microarray data. In Bioinformatics and Computational Biology Solutions Using R and Bioconductor. Statistics for Biology and Health. 2005.

Soneson, C. and Delorenzi, M. A comparison of methods for differential expression analysis of RNA-seq data. BMC bioinformatics 2013;14(1):91.

Subramanian, A., et al. Gene set enrichment analysis: a knowledge-based approach for interpreting genome-wide expression profiles. Proc Natl Acad Sci U S A 2005;102(43):15545–15550.

Tao, Y., et al. Information theory applied to the sparse gene ontology annotation network to predict novel gene function. Bioinformatics 2007;23(13):i529–538.

Tarazona, S., et al. Data quality aware analysis of differential expression in RNA-seq with NOISeq R/Bioc package. Nucleic acids research 2015;43(21):e140–e140.

Van Keymeulen, A., et al. Reactivation of multipotency by oncogenic PIK3CA induces breast tumour heterogeneity. Nature 2015;525(7567):119–123.

Vitali, F., et al. Developing a ‘personalome’for precision medicine: emerging methods that compute interpretable effect sizes from single-subject transcriptomes. Briefings in Bioinformatics 2017;20(3):789–805.

Wan, Y.W., Allen, G.I. and Liu, Z. TCGA2STAT: simple TCGA data access for integrated statistical analysis in R. Bioinformatics 2016;32(6):952–954.

Woolf, B. On estimating the relation between blood group and disease. Ann Hum Genet 1955;19(4):251–253.

Yang, X., et al. Single sample expression-anchored mechanisms predict survival in head and neck cancer. PLoS Comput Biol 2012;8(1):e1002350.

Zaim, S.R., et al. Evaluating single-subject study methods for personal transcriptomic interpretations to advance precision medicine. Bmc Medical Genomics 2019;12(5):96.

Zaim, S.R., et al. Emergence of pathway-level composite biomarkers from converging gene set signals of heterogeneous transcriptomic responses. Pac Symp Biocomput 2018;23:484–495.

